# Keeping students connected and engaged in a wet-lab research experience during a time of social distancing via mobile devices and video conferencing software

**DOI:** 10.1101/2021.09.30.462419

**Authors:** Michel Shamoon-Pour, Caitlin J. Light, Megan Fegley

## Abstract

Two major COVID-19 pandemic challenges presented for in-person instruction included adhering to social distancing guidelines and accommodating remote learners who were temporarily isolated or permanently participating from afar. At Binghamton University, our First-year Research Immersion (FRI) program was challenged with providing students with a wet lab course-based undergraduate research experience (CURE), an intense hands-on experience that emphasized student teamwork, lab protocol development, iteration, troubleshooting and other elements of the process of science that could not be replicated in a fully remote environment. We developed an innovative technology approach to maximize all students’ connection to the lab research experience utilizing dedicated mobile devices (iPod Touch) and video conferencing software (Zoom) to synchronously connect remote learners to in-person learners, peer mentors and instructors in our FRI research labs. In this way, despite limited lab capacities and fluctuating remote learning populations, we were able to connect remote learners to their peers and mentors in real-time and give them responsibilities that allowed them to be engaged and feel like meaningful participants in the research process. Although our students reported a preference for in-person labs, they noted this hybrid model was better than other traditionally employed remote-learning lab options. We believe lessons learned here can be applied to improve access to research in all situations and allow us to be prepared for other catastrophic disruptions to the educational system.

## Introduction

Due to the COVID-19 pandemic, social distancing and other safety measures have been particularly challenging in course-based undergraduate research experiences (CUREs). In order to achieve CURE pedagogical goals of students’ research competency (1), sense of project ownership (2) and self-identification with science and research communities (3), STEM CUREs rely on a range of in-person activities, namely lab work, field work, and face-to-face mentoring. There are also concerns about the adverse impact of moving to an online environment on students’ teamwork and communication skills (4,5). The disadvantages of abandoning in-person activities in CUREs necessitates that educators identify remote alternatives that can be quickly adopted in future catastrophic events (6).

In Binghamton University’s First-year Research Immersion (FRI) program, to comply with social distancing requirements, we implemented a technology-based solution for “wet lab” CUREs. FRI consists of 10 research tracks that provide first-year students with three sequential CUREs. In the first fall semester, students learn foundational research methods skills that are continually utilized as students progress through the CURE sequence. In the following spring semester, students learn discipline-specific research skills needed to complete their own research projects during the third and last CURE in the following fall semester (7). For this manuscript, we focused on the four tracks (Biofilms, Biogeochemistry, Biomedical Anthropology, Ecological Genetics) that in the second semester CURE (normally with two three-hour lab sessions per week) used the strategy described below on a regular basis in spring semester 2021.

While social distancing forced reduced lab capacity and in-person lab hours by 50%, our strategy utilized mobile devices and video conferencing software to maximize time engaged synchronously in research activities each week. Students were paired and rotated through in-person and remote roles. Overall, this technology allowed all students (in-person, remote choice for semester, or temporarily quarantined) to engage in the research environment synchronously (live), collaborate within teams and have meaningful responsibilities.

Although the intended audience of this report is primarily CURE educators, where wet lab or field work training is an essential part of students’ research and learning, the method can also be beneficial to synchronous remote teaching of any lab courses.

## Procedure

Zoom conferencing software on iPod Touch devices was used to connect remote students to the wet lab or field research experience. Thus, students interacted synchronously with their peers, undergraduate peer mentors (UGPMs), and research educators (REs) despite social distancing limitations (**Table 1**). In some cases, the technology allowed temporarily remote (quarantined) students to engage with an in-person lab partner, team, or UGPM to synchronously participate in wet-lab or field experiences. In other cases, the technology was used to permanently pair students in order to reduce the number of students in the lab. To optimize the utility of this strategy, in-person students were typically required to execute the physical lab or field work as in a traditional in-person CURE. Remote learners were typically responsible for things like directing and guiding their in-person lab partner through the experimental procedure and maintaining an electronic lab notebook (ELN). Together the in-person and remote students were to share and discuss their trouble-shooting, findings, and conclusions.

**Table 1.** Use of iPods for synchronous (live) remote lab learning.

| **Example uses of iPods for connecting remote learners** | **Paired lab partners in-person students + remote students** | **Remote students + UGPM** | **Remote students + in-person bench work student** | **Remote students + in the field students** | **Instructor demonstrations** |
| --- | --- | --- | --- | --- | --- |
| **Description** | One student in each pair attends lab in-person (A) while the other student attends remotely (B). Each week they rotate roles. | Remote learners (fully or temporarily) paired with an in-person UGPM in the lab. | Remote learners (fully or temporarily) paired with an in-person student or student team in the lab. | Remote learners (fully or temporarily) connected with UGPM, another student, or RE to participate in field work. | RE demonstrates detailed research skills live for in-person and remote learners while maintaining social distancing (doc-cam). |
| **Remote Responsibilities** | • Directing and guiding in-person lab partner through experimental procedure (calculations, explaining next steps, etc.)<br>• Maintaining ELN (recording procedure and observations, data collection, data analysis, etc.)<br>• Asking remote UGPMs clarifying questions | • Directing and guiding in-person UGPM through experimental procedure (calculations, explaining next steps, etc.) to demonstrate own understanding<br>• Maintaining ELN (recording procedure and observations, data collection, data analysis, etc.) | • Directing and guiding in-person lab partner/team through experimental procedure (calculations, explaining next steps, etc.) to demonstrate own understanding<br>• Maintaining ELN (recording procedure and observations, data collection, data analysis, etc.) | • Observation of samples and/or data in the field in realtime<br>• Maintaining ELN (recording procedure and field observations, data collection from field, data analysis, etc.) | • All students, in-person and remote, watch RE demo via Zoom<br>• Ask questions about demo<br>• Take notes on tips and tricks provided by RE |
| **In-person Responsibilities** | • Execution of lab techniques (protocols, instrumentation, etc.)<br>• Asking clarifying questions of RE or in-person UGPMs | • UGPM carrying out lab techniques (protocols, instrumentation, etc.) under direction of remote student(s)<br>• UGPM guides students through issues with series of counter-questions to encourage students to own experimental outcome | • Execution of wet lab techniques (protocols, instrumentation, etc.)<br>• Asking clarifying questions of RE or in-person UGPMs | • Travel to and collection of samples and/or data in field | • Same as remote responsibilities |
| **Typical Comments** | “Using iPods in the lab to communicate with and visualize my partner's work during lab is incredibly helpful. I feel as though this system works very well to connect lab partners working from different locations. I find maintaining my virtual lab notebook to be very helpful for organizing my data in a clear, accessible format that I can access at any time.” (FRI Student) | “I think that executing the experiment with the remote students over zoom was a good attempt as we were able to converse about what was happening in lab without their physical presence. They were also engaged throughout the process in asking questions about the steps.” (FRI UGPM) | “Even though I took classes online, my experience was not greatly affected. I was paired up with another student and he used an iPod to show me the whole process of each lab experiment. I also had the opportunity to direct him to conduct experiments according to the instructions I said, and then he would point out my mistakes and reminded me what steps can be done better.” (FRI Student) | “We made it outside in the beautiful weather and the hotspot worked for livestreaming the campus watershed tour.” (RE) | “Students seem to be appreciating the in-person /remote lab format and are really thinking about how to get the most out of it... Demonstrating how to load a gel is so much better with an iPod-eye view and I never want to demo it with people looking over my shoulder again.” (RE) |

Public health and lab safety were driving factors in purchasing iPod Touch devices. To avoid using students’ personal devices, iPods provided the most affordable, convenient, and adaptable technology with audio and video capabilities to facilitate mobile Zoom conferencing in the lab. Plastic cases and screen protectors were purchased to allow for proper cleaning and disinfection between use. The iPods met additional safety measures required for some of our labs (Biosafety Level 1 (BSL1), BLS2, Institutional Animal Care and Use Committee (IACUC)).

We asked for student feedback via google form twice throughout the semester to understand what was and wasn’t working for students. We also surveyed students about their remote learning experiences as part of our summative, end-of-semester, online Qualtrics survey. A complete list of survey questions is included in Appendix 1. Students also spoke to these interventions in their end-of-semester reflection essays (300-500 words) where they were asked to reflect on their professional and personal growth in the spring CURE. The study was approved by Binghamton University, IRB (no. STUDY00000104).

## Conclusion

Here we report student feedback outlining the challenges and success of using iPods and Zoom to link remote and in-person students in CURE lab environments.

### Formative assessment

Student responses to feedback surveys (survey #1, survey #2) during the semester revealed engagement issues for remote students due to: a) distractions during remote sessions and b) communication issues between in-person and remote students and between remote students and REs and UGPMs. In response, improvements included a) increased use of UGPMs for more intentional check-ins with remote students to help them stay connected and b) providing all students with more explicit in-person and remote student responsibilities. Although students indicated feeling less engaged during remote lab work compared to in-person lab work, a notable proportion of students who experienced in-person and remote work during the semester indicated feeling engaged “frequently” or “always” during remote lab work (49% survey #1, 58% survey #2).

### Summative assessment

In the end of semester assessment, the majority of all students indicated while conducting lab work remotely that they “frequently” or “always” felt they could connect and communicate in real time with their partner, UGPM, and/or RE (68%) and felt they were making meaningful contributions to the lab work and/or research project (62%). Considering the COVID-19 challenges and safety measures, student quotes from end of semester reflection essays (Table 1) indicated a positive student view of the iPod usage. When students were asked to rank their preferences for future lab experiences, an overwhelming majority of students indicated fully in-person lab experiences as their top preference (78%), followed by 50:50 Hybrid lab experience (9%), and Flipped lab experience (7%) (Fig. 1, with terms defined). Thus, the students recognized the value of the hybrid iPod method to enhance their experience, they also realized it did not replace in-person, hands-on scientific research experiences for them. A hybrid cannot provide as deep an immersion into research. For example, the level of iteration in a fully in-person wet lab research experience contributes markedly to building students’ professionalization and confidence (8).

**Figure 1.**
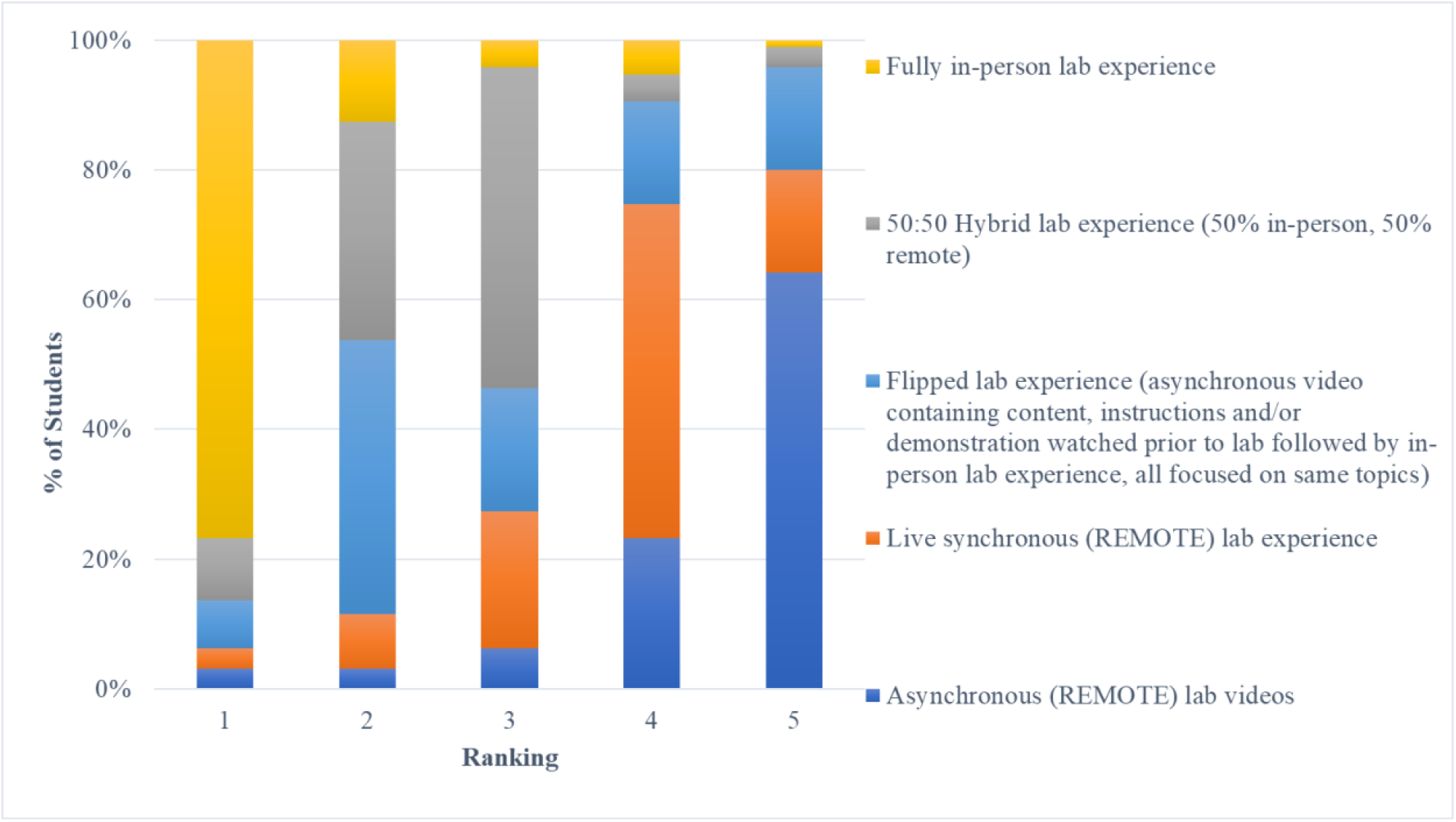
Student responses on the end of semester survey to the question “Based on all of your lab experiences, rank these lab learning methods according to your preferences for future lab experiences, with 1 being the method you would prefer the most,” n=95.

In order to keep remote learners engaged we made use of ELNs and utilized UGPMs remotely to jump through breakout rooms during lab to answer questions and ensure remote learners were on task, as they had been instructed as their responsibility, to guide their partners and document experiments in the ELN. Due to this method’s complex and dynamic technology use, we also compiled a series of recommendations and troubleshooting suggestions to help alleviate technology issues (Appendix 2).

Since teamwork and collaboration are central to CUREs and particularly important for students participating remotely, we used data collected from collaboration category questions in Corwin et al. Laboratory Course Assessment Survey (LCAS) (9). LCAS is a validated survey measuring students’ perceptions about their experience relative to categories of research-course design that science education research shows to relate closely to students’ engagement and retention in STEM majors. In terms of students’ perception of their exposure to collaborative activities, for our students, the mean for collaboration was above 95%, whereas the means for the reference data were below 90% of the range possible (**Table 2**). Even in a hybrid environment with reduced in-lab time and remote lab components, we were able to provide students with an environment conducive to building teamwork and collaboration skills.

**Table 2.** LCAS Collaboration Cluster Scores.

|  | **Mean** | **SD** | **n** | **% Range Possible** |
| --- | --- | --- | --- | --- |
| **Traditional students (LCAS ref)** | 20.87 | 4.0 | 68 | 87.0% |
| **CURE students (LCAS ref)** | 21.11 | 3.2 | 73 | 88.0% |
| **FRI Spring 2021 students** | 23.50 | 1.5 | 96 | 97.9% |
| Possible range of scores | 6–24 |  |  |  |

Although students prefer in-person lab experiences, our strategy can increase CURE and research access to students and educators beyond COVID-19. Students could benefit in future remote situations due to sickness, geography, or severe weather. When necessary, instrument and supply costs can be reduced due to lower numbers of in-person students. Institutions with limited financial resources and/or research instrumentation can collaborate with other institutions to give their students more research access through remote experiences. This method could also give educators, both K-12 and higher ed, the opportunity to receive remote training on CURE implementation and more.

## Acknowledgments

This study was made possible by resources provided by the Office of the Executive Vice President for Academic Affairs and Provost. We thank our FRI team, including REs, Jonathan Schmitkons and Christina Baer and numerous UGPMs for their technology implementation and discussion. We also thank Nancy Stamp for manuscript feedback and Binghamton’s IT department for technical assistance.

## Supplemental Materials

Appendix 1: Assessment Tools

Appendix 2: Technology guidance and recommendations to enhance remote student lab experience.

## Notes

### Competing Interest Statement

The authors have declared no competing interest.

