## Supplemental Materials for "Keeping students connected and engaged in a wet-lab research experience during a time of social distancing via mobile devices and video conferencing software"

**Appendix 1**: Assessment Tools

***Formative Assessment***

- **Feedback survey questions asked during the semester**
    - **In-Person Lab Work -** Questions should be answered based on your FRI course THIS SEMESTER.

Q1. When you are conducting in-person lab work for your FRI course this semester, do you feel engaged? Please give a brief reasoning for your response.

- Always
- Frequently
- Sometimes
- Rarely
- Never
- Not applicable (I have not conducted in-person lab work)

---

Q2. Have you conducted any FRI lab work remotely to fulfill your FRI lab requirements this semester? (Remote work could include attending lab via Zoom, analyzing data, computer programming, viewing a live experiment, entering information into an electronic lab notebook, working with an in-person lab partner, etc.)

- Yes
- No, I have not conducted FRI lab work remotely

---

- **Remote Lab Work -** Remote work could include attending lab via Zoom, analyzing data, computer programming, viewing a live experiment, entering information into an electronic lab notebook, working with an in-person lab partner, etc.

Q3. When you are conducting FRI lab work remotely, do you feel engaged? Please give a brief reasoning for your response.

- Always
- Frequently
- Sometimes
- Rarely
- Never
- Not applicable (I have not conducted lab work remotely)

---

Q4. While conducting FRI lab work remotely, do you feel like you can connect and communicate in real time with your partner, peer mentor and/or instructor? Please give a brief reasoning for your response.

- Always
- Frequently
- Sometimes
- Rarely
- Never
- Not applicable (I have not conducted lab work remotely)

---

Q5. While conducting FRI lab work remotely, do you feel like you are making meaningful contributions to the lab work and/or research project? Please give a brief reasoning for your response.

- Always
- Frequently
- Sometimes
- Rarely
- Never
- Not applicable (I have not conducted lab work remotely)

---

Q6. What works well when you are conducting FRI lab work remotely?

---

Q7. What would help increase your engagement and/or experience when conducting FRI lab work remotely?

---

***Summative Assessment***

- **Reflection Essay Prompt**
  Complete a reflection essay (XX points) by XXX. You must write 300-500 words typed discussing your professional & personal growth through the FRI research stream part I course. Upload your essay into your individual google drive folder as a word document. A periodic reflection essay is a valuable way for you to take stock of your professional and personal growth. So at the end of each FRI course we ask that you take some time to think and write about that in terms of this course.

- **End of Semester Survey Questions**
  - Part I. Learning Environment

Q1. I was familiar and comfortable with all the technology needed for the online portions of this course.

- Strongly agree
- Somewhat agree
- Neither agree nor disagree
- Somewhat disagree
- Strongly disagree

---

Q2. The following reduced my ability to participate/perform in the course:

| | Strongly agree | Somewhat agree | Neither agree nor disagree | Somewhat disagree | Strongly disagree |
|---|---|---|---|---|---|
| Access to technology | | | | | |
| Access or issues with internet | | | | | |
| Issues with technology | | | | | |
| Distractions from physical environment where I was online from | | | | | |
| Distractions from my personal technology (phone, computer, etc.) | | | | | |
| Health issues or stress (yours or family/roommate) | | | | | |
| Time zone differences | | | | | |
| Other | | | | | |

Q3. What portion of the live online synchronous lecture class time in the course was

| | 0 to 5% (rarely) | 6% to 20% (sometimes) | 21% to 40% (often) | 41% to 70% (most of the time) | 71% to 90% (almost always) | More than 90% (always) |
|---|---|---|---|---|---|---|
| Non-interactive lecture (mainly instructor talking) | | | | | | |
| Interactive Lecture (including polls and questions) | | | | | | |
| Sharing student comments and instructor responding to comments and questions | | | | | | |
| Small group work or discussion in breakout rooms | | | | | | |
| Individual work | | | | | | |

Q4. I felt comfortable asking questions in live online class.

- Strongly agree
- Somewhat agree
- Neither agree nor disagree
- Somewhat disagree
- Strongly disagree

Q5. I felt that the feedback I received in this course from the instructor, teaching assistants and peer mentors (including on assignments, during class, in virtual office hours, etc.) helped me to succeed.

- Strongly agree
- Somewhat agree
- Neither agree nor disagree
- Somewhat disagree
- Strongly disagree

---

Q6. Ease of my interactions with the instructor in this course experience compared to the average of my other college courses in similar disciplines was

- Much better
- Somewhat better
- About the same
- Somewhat worse
- Much worse

---

Q7. Ease of interacting with other students in this course experience compared to the average of my other college courses in similar disciplines was

- Much better
- Somewhat better
- About the same
- Somewhat worse
- Much worse

---

Q8. My overall educational experience in this course compared to the average of my other college courses in similar disciplines was

- Much better
- Somewhat better
- About the same
- Somewhat worse
- Much worse

---

Q9. When you were participating in or conducting the following activities in your FRI course this semester, did you feel **engaged**?

| | Always | Frequently | Sometimes | Rarely | Never | Not applicable |
|---|---|---|---|---|---|---|
| Attending live (synchronous) online lecture classes with instructor and other students present | | | | | | |
| Watching recorded content videos | | | | | | |
| Watching lab protocol videos | | | | | | |
| Reading of assigned texts (journal articles, etc.) | | | | | | |
| Completing graded assignments (lab notebook, problem sets, data analysis, experiments, etc.) | | | | | | |
| Completing writing assignments (technical report, proposal, etc.) | | | | | | |
| Team meetings | | | | | | |
| Office hours | | | | | | |
| Conducting in-person FRI lab work | | | | | | |
| Conducting FRI lab work remotely | | | | | | |
| Conducting FRI fieldwork | | | | | | |

Q10. What instructional strategies or course activities **contributed** most to your learning during **FRI LECTURE this semester**? Briefly explain why.

Q11. What instructional strategies or course activities **disengaged or discouraged** your learning during **FRI LECTURE this semester**? Briefly explain why.

- Part II. Laboratory Learning Environment

In this section of the survey, please respond to these questions in terms of your FRI laboratory experience this semester.

Q12. What instructional strategies or course activities **contributed** most to your learning during **IN-PERSON FRI LAB this semester**? Briefly explain why.

Q13. What instructional strategies or course activities **disengaged or discouraged** your learning during **IN-PERSON FRI LAB this semester**? Briefly explain why.

Q14. What instructional strategies or course activities **contributed** most to your learning during **REMOTE FRI LAB this semester**? Briefly explain why.

Q15. What instructional strategies or course activities <u>**dis**engaged or discouraged</u> your learning during <u>REMOTE</u> FRI <u>LAB</u> this semester? Briefly explain why.

---

Q16. While conducting <u>**FRI lab work remotely**</u> this semester, did you feel like you could connect and communicate in real time with your partner, peer mentor and/or instructor? AND give a brief reasoning for your response.

- Always ____________________________________________________
- Frequently ____________________________________________________
- Sometimes ____________________________________________________
- Rarely ____________________________________________________
- Never ____________________________________________________
- N/A (I have not conducted lab work remotely)

---

Q17. While conducting <u>**FRI lab work remotely**</u> this semester, did you feel like you were making meaningful contributions to the lab work and/or research project? AND give a brief reasoning for your response.

- Always ____________________________________________________
- Frequently ____________________________________________________
- Sometimes ____________________________________________________
- Rarely ____________________________________________________
- Never ____________________________________________________
- N/A (I have not conducted lab work remotely)

---

Q18. Which Binghamton University lab courses have you completed or are currently enrolled in?

---

Q19. My FRI lab experience this semester was _________ compared to my experiences in other **LAB** courses at Binghamton University AND give a brief reasoning for your response.

- Much better ____________________________________________________
- Somewhat better ____________________________________________________
- About the same ____________________________________________________
- Somewhat worse ____________________________________________________
- Much worse ____________________________________________________
- Not applicable (I have not completed or been enrolled in any other Binghamton courses which have a lab component.)

---

Q20. Based on all of your laboratory experiences, rank these laboratory learning methods according to your preferences for future laboratory experiences, with 1 being the method you would prefer the most: (Drag line of text to reorder ranking)
______ Asynchronous (REMOTE) laboratory videos (1)
______ Live synchronous (REMOTE) laboratory experience (4)
______ 50:50 Hybrid laboratory experience (50% in-person, 50% remote) (3)
______ Fully in-person laboratory experience (2)
______ Flipped laboratory experience (asynchronous video containing content, instructions and/or demonstration watched prior to lab followed by in-person lab experience, all focused on same topics) (5)

**Appendix 2**: Technology guidance and recommendations to enhance remote student lab experience.

1. Use one class Zoom link with breakout rooms for each student group or pair in order to allow RE/UGPMs to reach all students more efficiently.
2. Use of mobile hotspot to connect remote students for real-time fieldwork.
3. For best audio and noise reduction, in-person students use one wireless earbud.
4. Multifunctional iPod stands to allow for hands-free mobile or stationary work.
5. Designated help-space for remote learners staffed by RE/UGPMs (*i.e.,* Zoom breakout room).
6. IT department collaboration to confirm sufficient WIFI coverage and capacity in lab space. Institutions may have device management platforms available for mobile devices to configure and automate IT administration tasks.
